# Physics-informed multi-encoder adaptive optics enables rapid aberration correction for intravital microscopy of deep complex tissue

**DOI:** 10.64898/2026.03.07.710274

**Authors:** Xiangzhang Cheng, Bo Wang, Li Luo, Zhaowei Sun, Sicong He

## Abstract

Tissue-induced optical aberrations fundamentally constrain intravital microscopy in deep or complex biological specimens. While adaptive optics (AO) can compensate for these aberrations, conventional AO methods are limited by either guide-star dependency or slow correction speeds. Here, we develop MeNet-AO, a multi-encoder network-based AO method that enables rapid, guide-star-free aberration correction. By integrating a noise-resilient, structure-independent feature extraction model with a physics-informed multi-encoder architecture, MeNet-AO jointly decodes multiple large-amplitude aberration modes from wavefront-modulated image pairs, achieving an effective balance between prediction accuracy and temporal efficiency. Validated in living organisms, MeNet-AO improves fluorescence imaging in zebrafish brain and eye, enhances neuronal calcium transients and direction selectivity in mouse visual cortex, and enables subcellular-resolution microglial calcium imaging through thinned-skull windows – revealing spatiotemporally heterogeneous signaling patterns previously obscured by skull aberration. The speed and robustness of MeNet-AO in low-signal and scattering conditions establish it as a versatile platform for dynamic subcellular imaging deep within native tissue environments.

## Introduction

The orchestration of physiological functions in living organisms arises from precisely coordinated cellular dynamics within three-dimensional tissue microenvironments. *In vivo* observation of these spatiotemporal dynamics provides critical insights into cell functions and their roles in health and disease^1,2^. Two-photon fluorescence microscopy has emerged as an invaluable tool for intravital imaging due to its exceptional tissue penetration depth, subcellular resolution, inherent optical sectioning capability, and low phototoxicity^3–5^. However, the full potential of this technique for deep tissue imaging remains constrained by progressive degradation of point spread function (PSF) quality caused by specimen-induced optical aberrations. These wavefront distortions, arising from refractive index heterogeneities in biological specimens, imperfections in optical components, and system misalignments, impose fundamental limitations on achievable resolution and signal intensity at substantial depth, particularly when imaging dynamic processes in living organisms.

Adaptive optics (AO), originally developed for astronomical imaging, addresses these limitations and restores optimal imaging performance through wavefront correction using phase-modulating devices, such as deformable mirrors or spatial light modulators^6–9^. Current AO implementations predominantly employ either direct^10^ or indirect^11,12^ wavefront sensing strategies. Sensor-based approaches like Shack-Hartmann wavefront sensor (SHWS) enable rapid aberration measurement but require high-quality guide stars^13^. While early implementations utilized invasive injected fluorescent beads as artificial guide stars^14,15^, recent advancements leverage endogenous two-photon excited fluorescence signals as intrinsic guide stars to measure the tissue-induced aberrations and restore diffraction-limited resolution of two-photon microscopy^13,16–23^. Nevertheless, photon scattering and absorption in turbid biological tissues fundamentally limits the effectiveness of guide star-dependent methods at depth, particularly for intravital imaging applications where backscattered signal collection efficiency decreases exponentially with imaging depth^13,18,19^. On the other hand, indirect wavefront sensing methods circumvent this limitation by employing iterative optimization algorithms that maximize image sharpness or intensity metrics^24–29^, thereby eliminating the need for wavefront sensors. Though more adaptable to scattering specimen, these sensorless approaches suffer from slow convergence rates that render them unsuitable for imaging dynamic biological processes. Consequently, these fundamental limitations underscore the need for novel aberration correction strategies that combine the speed of direct wavefront sensing with the scattering robustness of indirect wavefront sensing methods.

The integration of deep learning with adaptive optics has opened new frontiers in computational microscopy, with convolutional neural networks demonstrating remarkable capabilities in image restoration tasks ranging from denoising to scattering compensation^30–41^. This synergy has driven the development of deep learning-based adaptive optics (DLAO) microscopy^42–45^, which aims to overcome the inherent compromises among temporal resolution, hardware complexity, and biological applicability in conventional AO implementations. Current DLAO methods can be broadly categorized into three paradigms: image translation^46,47^, wavefront prediction^48–53^, and hybrid approaches that simultaneously address both tasks^54,55^. The first paradigm employs end-to-end architectures to directly transform aberrated images into aberration-corrected outputs through supervised learning^46,47^. While eliminating the need for wavefront correctors enhances implementation simplicity, this approach fundamentally constitutes an ill-posed inverse problem – a single degraded observation may correspond to multiple aberration-corrected solutions^31,37,56–60^. Such inherent ambiguity raises concerns about biological fidelity when applied to dynamic intravital imaging where ground truth data is unavailable. The second paradigm adopts a physics-informed strategy where networks learn to map distorted image patterns to Zernike-mode aberration coefficients^48–53^ for subsequent hardware correction. Though theoretically more interpretable, these models require precise cross-domain mapping between spatial-frequency features and phase distortions – a relationship complicated by structural complexity in the samples. A third, hybrid paradigm has recently emerged to jointly estimate both the aberration-corrected images and the aberrated wavefront^54,55^. Nevertheless, this capability often necessitates retraining for each specific use case, placing significant demands on network tuning. As a result, most DLAO implementations thus remain restricted to simplified specimens with sparse structural features that facilitate aberration decomposition^48,49,52^. Recent innovations attempt to bridge this gap through a machine learning AO framework incorporating predefined wavefront modulations. By calculating PSF-related representations from modulated images and correlating them with Zernike modes^61–63^, such methods enhance generalization across biological specimens. A machine learning-based AO (MLAO) framework introduced by Hu et al. represents a pioneering effort that first demonstrated the feasibility of this strategy *in vivo* and extended it to multiple microscopy modalities^63^. However, its single-shot estimation accuracy is limited, and effective correction demands iterative refinement coupled with training on an extensive dataset. In addition, the requirement for sequential acquisition of multiple modulated images, which typically scales linearly with Zernike mode count, incurs substantial time costs when correcting high-order aberrations in thick tissues, presenting constraints for intravital imaging applications. These persistent challenges delineate four essential requirements for next-generation DLAO systems: (1) Concurrent prediction of low-order and high-order aberration modes with low latency (e.g. <5 seconds); (2) Robust compensation of large-amplitude aberrations (>0.5 μm root mean square (RMS)) prevalent in deep complex tissue environments; (3) Structure-independent generalization across diverse cellular morphologies and labeling patterns; (4) Noise-robust performance under photon-limited conditions. While existing methodologies have achieved advancements in addressing specific requirements, their combined implementation remains challenging for resolving dynamic subcellular processes in strongly scattering tissues at depths where large-amplitude, high-order aberrations emerge.

To address these challenges, we propose a Multi-Encoder Network-based Adaptive Optics (MeNet-AO) method for rapid, guide-star-free aberration correction in deep complex tissue imaging. Our approach introduces three key innovations: First, we develop a noise-robust, structure-independent feature extraction model that decouples aberration signatures from biological structures through systematic wavefront modulation and frequency-domain regularization. Second, through quantitative correlation analysis of Zernike mode modulation, we identify the optimal mode combination for efficient wavefront modulation and decoding, achieving an optimal balance between prediction accuracy (residual wavefront RMS <0.1 μm) and temporal efficiency (<5 seconds). Third, we architect a physics-informed multi-encoder network that simultaneously processes modulated image pairs through parallel feature extractors, enabling joint optimization of aberration mode decomposition with diverse amplitudes and orders. Experimental validation demonstrates MeNet-AO’s unique capability to resolve large-amplitude (up to 0.6 μm RMS) aberrations across seven Zernike modes (Z_5_-Z_11_) using only three modulated image pairs, completing full aberration correction within five seconds. We demonstrate MeNet-AO’s capabilities through intravital imaging of zebrafish nervous system and mouse cortex. Benchmarked against SHWS-based AO correction, MeNet-AO demonstrates equivalent aberration correction fidelity while eliminating guide star requirements. Notably, MeNet-AO demonstrates unique advantages in low-signal and highly scattering environments where conventional SHWS-based AO fails due to guide star degradation. In mouse visual cortex, MeNet-AO enhanced two-photon imaging precision to resolve visual-evoked calcium transients with improved neuronal classification fidelity, enabling functional decoding of stimulus-specific neural responses. Crucially, through thinned-skull preparations and MeNet-AO correction, we achieved minimally invasive, subcellular-resolution recording of microglial calcium dynamics. We captured propagating calcium waves along submicron processes and revealing compartmentalized calcium heterogeneity in distal branches, while circumventing microglial activation artifacts inherent to open-skull methods. These validations establish MeNet-AO as an effective platform for high-resolution and high-fidelity investigation of neuronal and glial dynamics in native microenvironments. MeNet-AO’s unique capacity to rapidly correct high-order aberrations at tissue-prevalent amplitudes enables unprecedented access to dynamic subcellular processes in deep complex tissues.

## Results

### Fast and structure-independent aberration estimation via multi-encoder networks

Fluorescence images inherently encode specimen-induced wavefront aberrations through intensity distortions. However, direct aberration estimation from raw images remains ill-posed due to structural complexity and labeling variability. In order to extract aberration-relevant features independent of the object pattern, we integrated three processing modules: (1) phase modulation for aberration signature encoding, (2) noise-robust feature extraction through Fourier-PSF transformation, and (3) multi-encoder neural network for parallelized mode decomposition. The framework of the MeNet-AO method is shown in Fig. 1a. First, predefined aberrations were applied to the raw aberrated images as wavefront modulations. By applying phase shifts with opposing signs to three selected Zernike modes, we generated modulated image pairs, preserving intrinsic specimen aberrations while encoding aberration responses. Through Fourier-domain calculation of the modulated images with appropriate regularization, we derived PSF-like representations that decouple aberrations from specimen structures and exhibit improved robustness to large aberrations and noise (Supplementary Note 1-2, Materials and Methods). Innovatively, we developed a multi-encoder neural network that comprehensively leverages distorted content differences following various modulations to accurately decode multiple aberration modes from the images and reconstruct the wavefront (Fig. 1a, Supplementary Fig.1). Each encoder was trained to extract spatial features from wavefront modulation of specific Zernike modes to provide a diverse representation of the input. At last, a feature coalescing module adaptively synthesizes these representations into comprehensive wavefront predictions as Zernike coefficients. This multi-encoder framework effectively extracts and combines features from specific modulations, enhancing aberration estimation and maximizing modulation efficiency. In contrast to conventional modal-based sensorless AO methods, which require time-consuming iterative trial-and-evaluate processes for each aberration mode, MeNet-AO achieves multi-mode estimation through a single forward pass, reducing the correction time from typically over 30 seconds to less than 5 seconds.

**Fig. 1.**
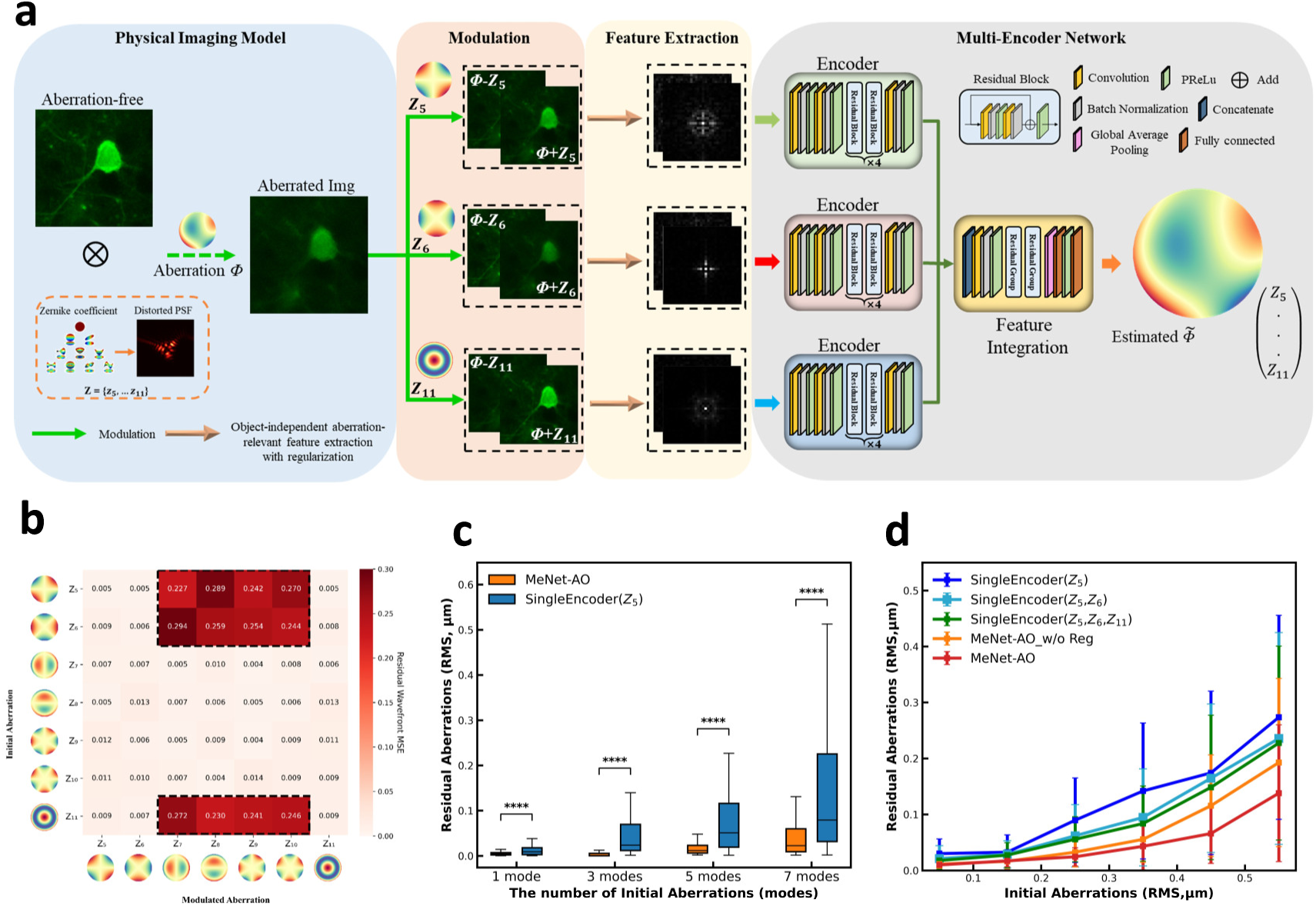
MeNet-AO schematic and simulation validation. **(a)** Schematic of the MeNet-AO workflow. The aberrated images are modulated with conjugate pairs of three centro-symmetric Zernike modes (Z_5_, Z_6_, Z_11_), followed by specialized feature extraction to obtain aberration-related features. These features are fed into a neural network comprising three expert encoders and a feature integration module, which outputs the predicted wavefront in terms of Zernike modes. **(b)** Cross-modulation correlations between different aberration modes. The amplitude of initial aberrations ranged from -0.5 μm to 0.5 μm (root-mean-square, RMS), with all other parameters kept constant except for the initial and modulated aberrations types. **(c)** Residual RMS wavefront error comparison between MeNet-AO (three-mode modulation with regularization) and SingleEncoder (modulated by oblique astigmatism, Z_5_, without regularization) (Materials and Methods) correction with increasing aberration complexity. The initial wavefront distortions range from 0 to 0.6 μm (RMS, n=100). The x-axis labels (1 mode, 3 modes, 5 modes, and 7 modes) correspond to cases with one (Z_5_), three (Z_5_-Z_7_), five (Z_5_-Z_9_), and seven (Z_5_-Z_11_) initial aberration modes, respectively. Two-sided paired t-test: ****p < 0.0001. **(d)** Residual RMS wavefront error versus initial aberration magnitude (0-0.6 μm RMS, seven Zernike modes (Z_5_-Z_11_), n=50) for different prediction models and modulation strategies. Blue curve: single-encoder model (Materials and Methods) utilizing stacked inputs modulated with conjugate pairs of Z_5_ mode, without regularization; Cyan curve: single-encoder model utilizing stacked inputs modulated with conjugate pairs of Z_5_ and Z_6_ modes, without regularization; Green curve: single-encoder model utilizing stacked inputs modulated with conjugate pairs of Z_5_, Z_6_ and Z_11_ modes, without regularization; Orange curve: MeNet model utilizing parallel inputs modulated with conjugate pairs of Z_5_, Z_6_ and Z_11_ modes, without regularization; Red curve: MeNet model utilizing parallel inputs modulated with conjugate pairs of Z_5_, Z_6_ and Z_11_ modes, with regularization. The x-axis is divided into bins of 0-0.1 μm, 0.1-0.2 μm, 0.2-0.3 μm, 0.3-0.4 μm, 0.4-0.5 μm, and 0.5-0.6 μm RMS.

Next, we characterized the performance of this multi-encoder model using image datasets with simulated aberrations. Specifically, fluorescence images were acquired from fixed brain slices with GCaMP-labeled neurons and DAPI-labeled cell nuclei. Then the images were degraded sequentially by simulated initial aberrations (Z_5_-Z_11_, Noll index, 0-0.6 μm RMS), modulated phase patterns (±0.2 μm amplitude on selected Zernike modes), and noise (Poisson-Gaussian mixture, Supplementary Note 3). The network was trained to map feature representations extracted from modulated images to the ground-truth Zernike coefficients, establishing a physics-constrained learning framework (Supplementary Fig. 2).

Recent studies revealed that certain Zernike mode, such as astigmatism, has better capability to decode other aberration modes when used as modulation^63,64^. To determine the optimal aberration types for wavefront modulation, firstly we calibrated the cross-modulation correlations between different aberration modes. Fluorescence images were aberrated by initial aberrations of individual Zernike mode and were then modulated by each aberration mode (Z_5_-Z_11_). A single-encoder model was used to estimate the initial wavefront aberration. Through predictive coupling analysis of Zernike modes, we quantified each mode’s cross-predictive capacity in aberration estimation (Fig. 1b). The results identify optimal mode combinations that maximize prediction power while minimizing redundant modulation. Notably, we found that the centro-symmetric modes (Z_5_ oblique astigmatism, Z_6_ vertical astigmatism, Z_11_ primary spherical aberration) demonstrated superior multi-modal predictability. In contrast, non-centrosymmetric modes (Z_7_ vertical coma, Z_8_ horizontal coma, Z_9_ vertical trefoil, Z_10_ oblique trefoil) showed limited cross-modal utility^63,64^ (Fig. 1b, Supplementary Fig. 3), indicating that coma and trefoil are suboptimal choices for modulation. This mechanistic insight inspired our strategy to select Z_5_, Z_6_, and Z_11_ as modulation modes for maximal cross-modal predictivity. This also motivated us to develop specialized encoders tailored to different modulation modes, aiming to extract aberration-related features with the highest efficiency.

Then we quantitatively compared the aberration estimation capabilities of multi-encoder versus single-encoder schemes across varying Zernike orders and amplitudes (Fig. 1c-d, Supplementary Fig. 4a-b). Notably, our multi-encoder approach utilizing three optimized centro-symmetric aberration modes (Z_5_, Z_6_, Z_11_) outperformed the single-encoder configuration with stacked inputs of the same modes in predicting aberrations comprising one to seven Zernike modes, exhibiting robust correction across varying aberration complexities (Fig. 1c, Supplementary Fig. 4a-b). Moreover, MeNet-AO demonstrates superior prediction accuracy compared to the single-encoder especially for large aberrations (>0.3 μm RMS). Results show that MeNet-AO maintained ∼0.1 µm accuracy (close to the diffraction limit^65,66^) for seven-mode aberrations with RMS wavefront errors up to 0.6 µm (Fig. 1d; Supplementary Fig. 4d). The results demonstrate MeNet-AO’s capability to accurately predict high-order, large-amplitude aberrations, which is an essential requirement for effective wavefront correction in intravital deep tissue imaging applications. In addition, to enhance noise robustness, we implemented a frequency-domain regularization strategy for high-efficiency feature extraction (Supplementary Note 2). Evaluation on both synthesized image data (Supplementary Note 3, Supplementary Fig. 5) and simulated aberration data (Fig. 1c, d, Supplementary Fig. 4a-d) consistently show that frequency-domain regularization further enhances the aberration prediction accuracy of MeNet-AO. Furthermore, we performed a systematic comparison with MLAO. The results demonstrate that, with only a single correction cycle, MeNet-AO delivers markedly superior performance in handling complex, large-amplitude aberrations (Supplementary Fig. 6a, b). Meanwhile, the magnitude of the modulated modes (Supplementary Fig. 4e) and the network architecture (Supplementary Fig. 7) were also optimized for MeNet-AO.

### MeNet-based computational adaptive optics microscopy

To validate the practical performance of MeNet-AO, we implemented an adaptive-optics two-photon microscopy system incorporating (1) a deformable mirror (DM) for both wavefront modulation and correction, (2) a Shack-Hartmann wavefront sensor (SHWS) for wavefront measurement as reference, and (3) a high-performance workstation hosting the MeNet-AO framework (Fig. 2a, Supplementary Fig. 8). In the MeNet-AO workflow, the DM is applied to introduce paired modulation aberrations to generate PSF-discriminative image pairs. These modulated images are then processed by MeNet to predict wavefront aberrations, which are subsequently compensated by the DM. For controlled validation, we corrected the optical system aberrations in advance, and then introduced predefined aberrations to aberration-free slice samples using the DM, enabling quantitative assessment of MeNet-AO’s prediction accuracy. The direct wavefront sensing (DWS) module provides reference aberration measurements using two-photon-excited fluorescent guide stars. Comparative analysis against DWS-corrected images establishes rigorous performance benchmarks for MeNet-AO.

**Fig. 2.**
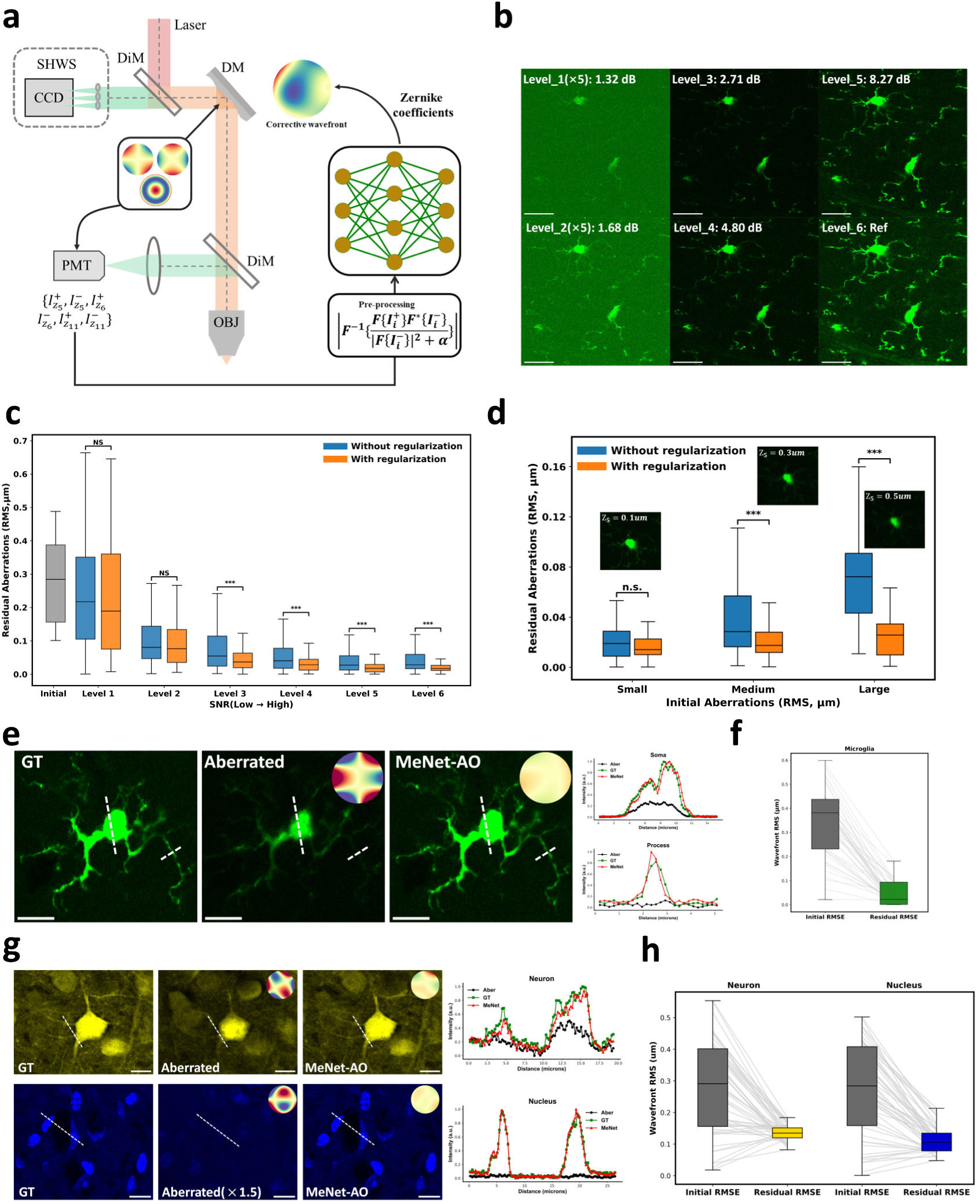
MeNet-AO microscopy system integration and validation. **(a)** Schematic diagram of two-photon fluorescence microscope integrating MeNet-AO framework for real-time aberration evaluation and correction. DM: deformable mirror; SHWS: Shack-Hartmann wavefront sensor; DiM: dichroic mirror; OBJ: objective lens; PMT: photomultiplier tube. The SHWS-based direct wavefront sensing module serves as the benchmark for MeNet-AO validation in intravital imaging. **(b)** Two-photon fluorescence images of brain slices (green: GFP-labeled microglia) acquired under varying excitation laser powers. The image datasets with different signal-to-noise ratios (SNR) were used to test the noise robustness of MeNet-AO for aberration estimation. For better visualization, the raw signals of the low-SNR images (Level_1 and Level_2) were digitally multiplied by 5, as indicated. **(c)** Residual RMS wavefront error versus SNR levels for the model with/without regularization. The model was trained on a dataset of 1,200 images (200 per SNR level), and tested on 100 images per SNR level. Oblique astigmatism (Z_5_) served as both the initial and modulation aberration, with the initial aberration amplitudes uniformly sampled from 0 to 0.5 μm RMS. Two-sided paired t-test: ***p < 0.001; NS, not significant. **(d)** Residual RMS wavefront error versus initial aberration amplitude for MeNet-AO with/without regularization. Small: 0-0.2 μm RMS; Medium: 0.2-0.4 μm RMS; Large: > 0.4 μm RMS. Two-sided paired t-test: ***p < 0.001; NS, not significant. **(e)** Validation of MeNet-AO on GFP-microglia slices. Representative images: ground truth (GT, system aberration corrected), aberrated (degraded by DM-induced aberrations), and MeNet-AO corrected. The MeNet-AO model was trained on 5000 images (initial aberrations: seven modes Z_5_-Z_11_; modulation modes: Z_5_, Z_6_, and Z_11_). Line profiles measured along the white dashed lines demonstrate that MeNet-AO restores imaging quality comparable to the GT. **(f)** Statistical analysis of MeNet-AO correction for microglia images within the same field of view (n=50; initial aberration amplitudes: 0-0.6 μm RMS). **(g)** Cross-sample validation: tdTomato-neuron and DAPI-nuclei slice images corrected by microglia-trained MeNet-AO model. Representative images with corresponding line profiles showing the restoration of image quality by MeNet-AO. For better visualization, the raw DAPI-nuclei slice image was digitally multiplied by 1.5, as indicated. **(h)** Statistical analysis of MeNet-AO correction for neuronal and nuclei images (n=50; initial aberration amplitudes: 0-0.6 μm RMS). Neuron and nuclei images were acquired from separate fields of view. Scale bar, 20 μm (b), 10 μm (e,g).

While the simulated datasets incorporated microscopy parameters (excitation wavelength, numerical aperture), discrepancies between simulated and experimental aberrations remained, arising from DM actuator resolution limits and system misalignment errors. As evidenced by prior studies, networks trained on experimental datasets outperform simulation-only counterparts^31,35,37,40,60^. We therefore collected ground-truth datasets by introducing mixed aberrations (Z_5_-Z_11_, 0-0.6 μm RMS) via DM to aberration-free, fixed-tissue slices. The resulting modulated image pairs with known aberration targets enabled effective training transferable to intravital imaging scenarios. Next, we systematically assessed MeNet-AO’s performance across regularization parameters and identified the optimal value that yielded the best performance (Supplementary Fig. 9a). The powerful feature extraction capability of the multi-encoder architecture reduced data dependency, achieving high-fidelity prediction (residual wavefront <0.05 μm RMS) with minimal specimen requirements. Remarkably, a training dataset of 5000 modulations acquired from only five regions within a single tissue slice was sufficient to deliver robust correction performance (Supplementary Fig. 9b).

Validation under controlled signal-to-noise ratio (SNR) conditions demonstrated the indispensable role of regularization for noise-resilient aberration extraction (Fig. 2b-c, Supplementary Figs. 10 and 11). Additionally, regularization also enhanced MeNet-AO’s ability to estimate large-amplitude aberrations, improving accuracy threefold for aberrations exceeding 0.4 μm RMS (Fig. 2d, Supplementary Fig. 11). On experimentally acquired datasets with varying SNRs and aberration amplitudes, MeNet-AO outperforms MLAO, demonstrating superior robustness to noise and complex aberrations (Supplementary Fig. 6c, d). Furthermore, to address the practical challenge of spatial shifts between modulated frames during live imaging, we implemented a shift-correction strategy that preserves MeNet-AO’s efficacy under image translation (Supplementary Note 4, Supplementary Fig. 12).

We further assessed MeNet-AO’s cross-structural generalization capability using a mouse brain slice with triple-color labeling identifying microglia (CX3CR1-GFP), neurons (Thy1-tdTomato) and nuclei (DAPI) within identical anatomical regions (Fig. 2e-h, Supplementary Fig. 13). This co-registered dataset enabled direct evaluation of aberration prediction consistency across fundamentally distinct cellular structures under identical optical conditions. The results show that, when trained exclusively on GFP-labeled glial images (Fig. 2e, f), MeNet-AO maintained high prediction accuracy when tested on structurally distinct nuclei and neuron images (Fig. 2g, h). Notably, the model exhibited robust transferability, generalizing effectively between any two image patterns (Supplementary Fig. 13). This structure-independent aberration correction performance originates from our physics-informed feature-disentangling preprocessing strategy, which decouples aberration information from image structures prior to network input. Collectively, MeNet-AO’s cross-sample generalization, robustness to noise, and ability to predict aberrations of large amplitude and high complexity provide a strong foundation for its application to intravital imaging of deep complex tissues.

### MeNet-AO enables high-resolution *in vivo* imaging of zebrafish larvae

To validate MeNet-AO’s generalizability from tissue slices to *in vivo* contexts, we deployed the MeNet model trained on DM-induced aberrations in fixed slices for live imaging of *Tg(Neurod:GFP; flk1:DsRedx)* transgenic zebrafish, in which neurons and vessels were labeled with GFP and DsRedx, respectively. Zebrafish larvae at 4 days post-fertilization (4 dpf) were dorsally mounted under anesthesia for live imaging. (Fig. 3a). We evaluated MeNet-AO correction at various locations within the zebrafish (Fig. 3b, Supplementary Fig. 14a). At a depth of 175 µm in the brain, image resolution and contrast were degraded by aberrations arising from brain curvature and refractive index heterogeneity (Fig. 3c). Following aberration measurement and correction by MeNet-AO, the two-photon fluorescence signals from neuronal somata and axons increased by one-to two-fold (Fig. 3c, Supplementary Fig. 14b). Additionally, frequency domain analysis revealed that finer details, indicated by high-frequency components, were recovered after MeNet-AO correction (Fig. 3c).

**Fig. 3.**
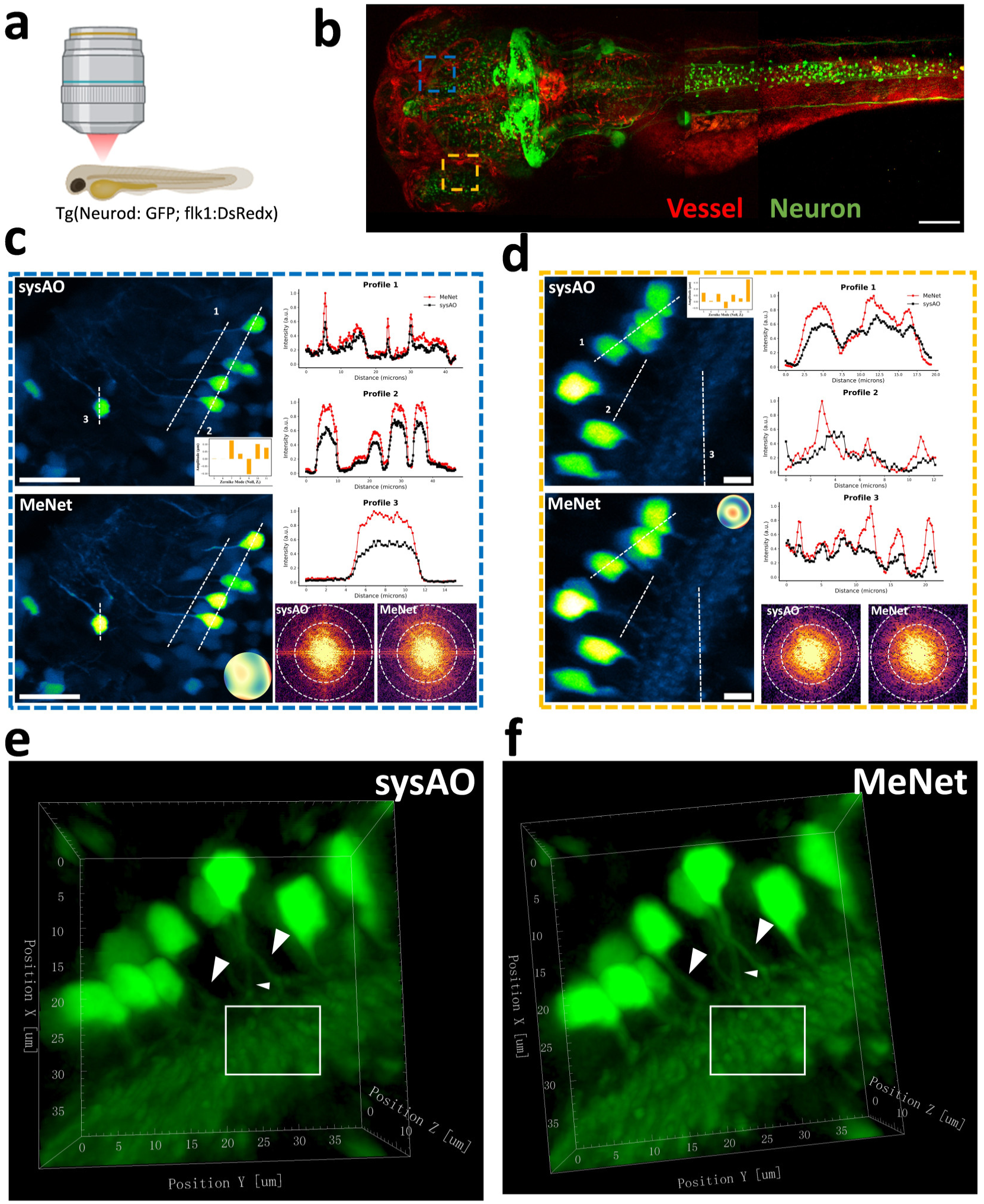
MeNet-AO enhances *in vivo* neuronal imaging in zebrafish. **(a)** Experimental setup for in vivo imaging of zebrafish larvae. **(b)** Two-color fluorescence image (maximum intensity projection) of *Tg(Neurod:GFP; flk1:DsRedx)* zebrafish larvae at 4 days post-fertilization (dpf). Green: neurons; Red: blood vessels. **(c)** Maximum intensity projections of hindbrain neurons (172–178 μm depth) with system AO and MeNet-AO correction. Insets display the corresponding corrective Zernike modes and reconstructed wavefront. The line profiles along the white dashed lines (normalized intensity) and the frequency representations of the images show the improved signal intensity and restored details after MeNet-AO correction. **(d)** Maximum intensity projections of retinal neurons (132–144 μm depth) with system AO and MeNet-AO correction. The annotations and layout are the same as in (c). **(e,f)** 3D reconstruction (39×39×13 𝜇𝑚^ଷ^) of GFP-labeled retinal neurons with system AO and MeNet-AO correction. MeNet-AO enhances visualization of dense neuropils within the inner plexiform layer (white boxed region) and neuronal axons (arrowheads) in the zebrafish retina. Scale bar, 100 μm (b), 20 μm (c), 5 μm (d).

Next, we targeted the zebrafish eye, where pronounced curvature induced severe aberrations that diminished fluorescence signals from neuronal somata and axons of retina (Fig. 3d). MeNet-AO correction enhanced retinal neuron signals in the inner nuclear layer (INL) while clearly resolving their axons projecting to the inner plexiform layer (IPL) (Fig. 3d, Supplementary Fig. 14c). 3D reconstructions demonstrated enhanced clarity of the dense neuropil within the IPL and clearly delineated neuronal axons following MeNet-AO correction (Fig. 3e,f).

In contrast, sensor-based AO produced suboptimal corrections at identical locations, due to degraded and uneven spot patterns resulting from endogenous absorption within tissues (Supplementary Fig. 14e). This limitation persisted across multiple locations, depths, and orientations (lateral or dorsal mounting, Supplementary Fig. 14g, 15c), reflecting the constraints of guide-star-based wavefront sensing approaches in complex tissues.

These results demonstrate that the slice-trained MeNet model can be directly transferred to diverse *in vivo* contexts. This further validates MeNet-AO’s ability to generalize across different fluorescent labeling patterns and complex tissue geometries in live organisms.

### MeNet-AO enhances *in vivo* structural and functional imaging in mouse cortex through open-skull cranial windows

*In vivo* recording of neuronal activity within natural physiological contexts is crucial for functional characterization and classification of cortical neuron types. However, intrinsic tissue aberrations and cranial window aberrations degrade spatial resolution and contrast at depth, critically compromising the fidelity of structural and functional neuronal imaging *in vivo*. To validate MeNet-AO for cortical imaging, we performed *in vivo* imaging of GCaMP6f-expressing neurons in the visual cortex of Thy1-GCaMP6f mouse through open-skull cranial windows (Fig. 4a). Structural imaging at 170 μm depth revealed blurred soma contours and indistinct neurites without aberration correction (Fig. 4b), attributed to tissue-induced wavefront distortions. Following MeNet-AO wavefront measurement and correction using the GCaMP6f fluorescence signal, images exhibited sharply defined soma boundaries and clear dendritic structures. Line profile analysis across identical neuronal dendrites confirmed substantial improvements in signal intensity and resolution after AO correction (Fig. 4b).

**Fig. 4.**
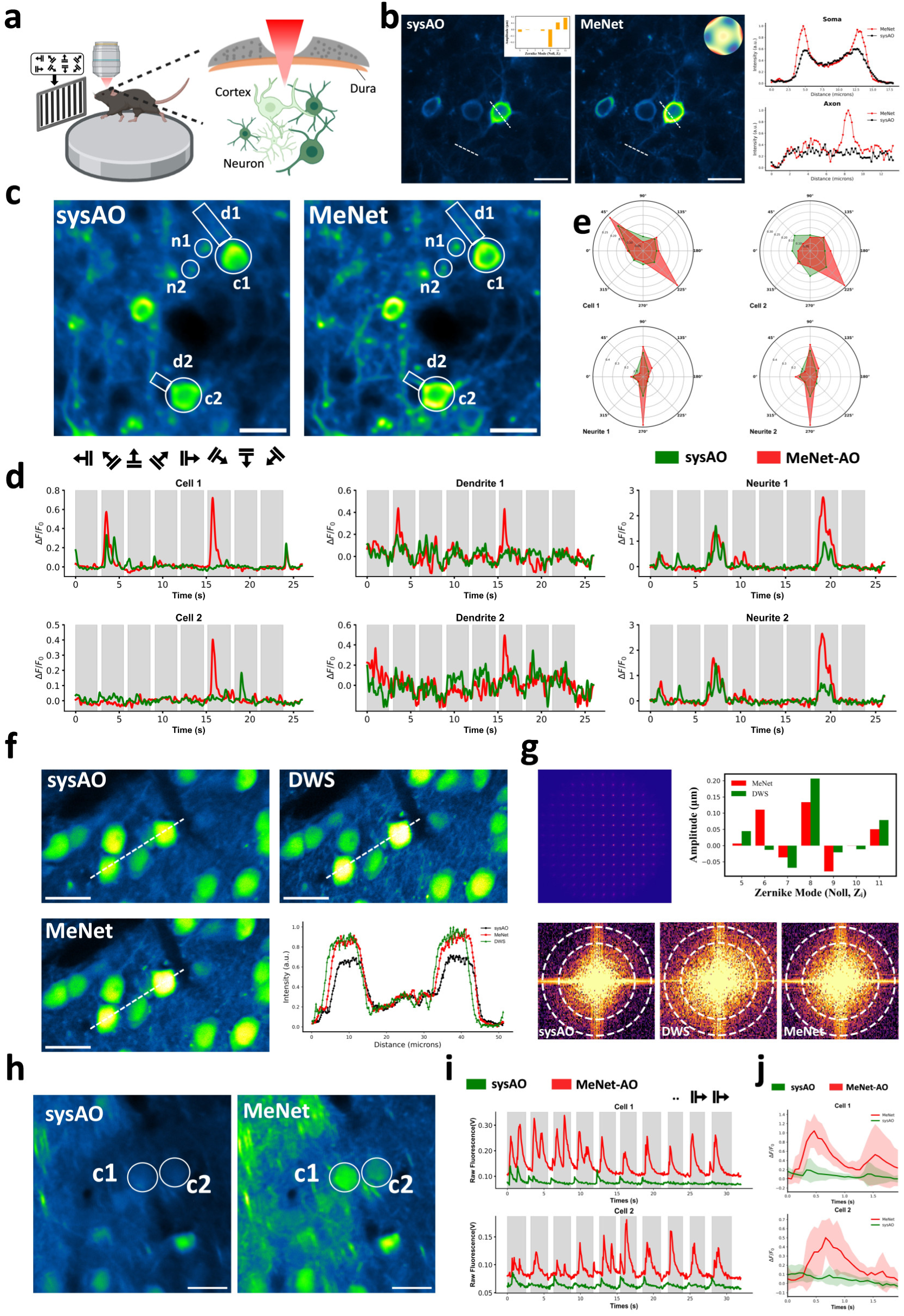
MeNet-AO enhances structural and functional neuronal imaging in mouse cortex. **(a)** Experimental schematic of imaging the mouse visual cortex through an open-skull cranial window during visual stimulation. **(b)** Representative GCaMP6f-labeled neuronal images at 170 μm below pia, with system AO correction (left) and MeNet-AO correction (right). Dashed lines indicate the positions of intensity profile comparison. The images are average projections of 20 frames. **(c)** Maximum intensity projection images of GCaMP6f-labeled neurons (26 seconds recording, one stimulus cycle) at 230 μm depth, with system AO correction (left) and MeNet-AO correction (right). **(d)** Calcium transients (smoothed) from the soma/dendrites/neurites outlined in (c) during visual stimulation at eight orientations, with system AO correction (green) and MeNet-AO correction (red). Each curve represents the average response across six stimulus repetitions. **(e)** Orientation tuning maps of Cell 1, Cell 2, Neurite 1 and Neurite 2 with system AO correction (green) and MeNet-AO correction (red), revealing enhanced direction selectivity. **(f)** Average intensity projection images of NLS-Dendra2- and jGCaMP8s-labeled neurons with system AO, MeNet-AO, and DWS-AO correction. Line profiles along the white dashed lines show comparable image quality between MeNet-AO and DWS-AO corrections. **(g)** Top left: A high-quality spot pattern acquired from bright guide stars using Dendra2 fluorescence, enabling effective wavefront measurement via DWS. Top right: Zernike mode coefficients reconstructed by MeNet and DWS. Bottom: Frequency representations of images highlight restoration of high-frequency details after MeNet-AO and DWS-AO correction. **(h)** Maximum intensity projection images of Dendra2/jGCaMP8s-labeled neurons recorded over 40 seconds at 245 μm depth, with system AO correction (left) and MeNet-AO correction (right). **(i)** Calcium signal traces from Cell 1 and Cell 2 outlined in (h), in response to repeated visual stimulation with drifting gratings in the eastward direction. **(j)** Averaged calcium transients (ΔF/F₀) of Cell 1 and Cell 2 with system AO correction (green) and MeNet-AO correction (red). Dark curves: averaged calcium transients. Shaded areas: the standard deviations of the calcium transients for each visual stimulus. Scale bar, 20 μm (b,c,f,h).

To assess neuronal function in the visual cortex, we recorded calcium dynamics of neurons in response to drifting grating visual stimuli. In densely labeled Thy1-GCaMP6f mouse cortex, calcium transients from neuronal somata and dendrites are frequently contaminated by surrounding neuropil signals^67^. This artifact is further amplified as tissue aberrations expand the laser’s focal volume, underscoring the need to separate neuropil signals from somatic activity in intravital calcium imaging. To evaluate MeNet-AO’s performance in functional imaging, we conducted high-resolution two-photon calcium imaging of neuronal dendrites and somata in layer 2/3 of mouse primary visual cortex (V1) at 230 μm depth. Visual stimuli, consisting of gratings drifting in eight different directions, were presented to the animals while time-lapse images were acquired sequentially with and without AO correction (see Section Materials and Methods). Consistent with structural imaging results, MeNet-AO enhanced the brightness of somata, dendrites, and spines in the averaged time-lapse image (Fig. 4c). Following motion correction and segmentation, we analyzed calcium transients in representative orientation-selective neurons and found that MeNet-AO correction not only significantly enhanced response amplitudes at each cell’s preferred grating orientation (c1, d1, n1, n2) but also revealed distinct direction selectivity (c2, d2) (Fig. 4d&e, Supplementary Fig. 16a), which is more variable than orientation preference^68,69^. Importantly, direction-tuning properties were preserved in corresponding dendritic segments (d1 & d2) following AO correction (Fig. 4c-e). Meanwhile, MeNet-AO enabled the precise localization of fine neurites and characterization of their orientation selectivity with improved fidelity (Supplementary Fig. 16b-d), leveraging AO-enhanced contrast and resolution. Two representative neurites (n1 & n2) encoded sufficient direction information and exhibited stronger visually evoked responses to their preferred stimulus direction than adjacent somata (Fig. 4c-e). This aligns with prior findings that the axo-dendritic processes and neuropil microdomains in V1 show greater reliability, higher spatial coherence, and similar decoding accuracy in direction discrimination compared to somatic activity^70^. Notably, the GCaMP6f signal is too weak for the wavefront sensor to accurately measure aberrations (Supplementary Fig. 16e), highlighting the limitation of the DWS approach under low SNR conditions and underscoring the necessity of MeNet-AO for intravital imaging. These results demonstrate MeNet-AO’s capacity to resolve the functional organization of cortical circuits at subcellular scales, revealing high-fidelity correlations between somatic and neuritic calcium dynamics previously obscured by aberrations.

To benchmark MeNet-AO against DWS-AO *in vivo*, we employed viral co-expression of nls-Dendra2 and jGCaMP8s in mouse visual cortex. This strategy generated bright nuclear-localized fluorescence (Dendra2) for guide-star-based aberration measurement, overcoming insufficient signal from Thy1-GCaMP6f-labeled neurons. At 130 µm depth, using the bright Dendra2 fluorescence as the measurement signal, DWS-AO and MeNet-AO achieved comparable correction on neurons, both enhancing resolution and contrast to resolve fine dendritic structures (Fig. 4f). Both MeNet and DWS produced similar aberration measurement results, with astigmatism and coma emerging as the dominant Zernike modes (Fig. 4g). Frequency-domain analysis revealed that high-frequency components, indicative of finer structural details, were recovered after both MeNet-AO and DWS correction (Fig. 4g). Functional imaging also reveals heterogeneous calcium dynamics in response to visual stimulation with different drifting directions (Supplementary Fig. 17). However, under the more challenging condition of greater imaging depth (245 µm) and low fluorescence intensity, DWS-AO failed because of guide star degradation (Supplementary Fig. 17d), whereas MeNet-AO still maintained robust performance owing to its noise-resilient, structure-independent design (Fig. 4h). Subsequently, we quantified functional imaging enhancement at this depth by recording calcium transients during ten consecutive visual stimuli of the same orientation, with and without aberration correction. Results show that MeNet-AO correction increased baseline fluorescence (F_0_) in neuronal traces by over twofold and calcium response amplitudes (ΔF/F_0_) by over fivefold (Fig. 4i). The trial-averaged signals highlighted the enhanced calcium traces after MeNet-AO correction (Fig. 4j). Critically, MeNet-AO correction in deep cortex enabled accurate decoding of neuronal responses to repeated identical stimuli, which is otherwise undetectable without correction. Furthermore, we compared MeNet-AO directly with system AO and MLAO at multiple imaging depths in the mouse brain. When limited to a single correction cycle, MLAO’s performance degraded with depth, while MeNet-AO remained effective across all depths tested (Supplementary Fig. 18), underscoring MeNet-AO’s superior efficacy in deep tissues where complex aberrations and low signal prevail. These results demonstrate MeNet-AO as a versatile solution for intravital imaging across diverse labeling strategies and SNR conditions, outperforming DWS-AO in deep tissues while eliminating guide star dependency.

### MeNet enables minimally invasive structural and functional imaging of cortical microglia via thinned-skull windows

Microglia, the resident immune cells in the central nervous system, are highly dynamic and sensitive to their surrounding physiological environment. While open-skull cranial windows provide optical access for brain imaging, they inevitably induce inflammation and activate microglia^71,72^. Therefore, minimally-invasive, high-resolution observation of microglial structure and function is crucial for accurately studying brain physiopathological states. To avoid microglial activation, we performed *in vivo* imaging of mouse brains through thinned-skull preparations (Fig. 5a, Section Materials and Methods), but the inherent refractive index heterogeneity and severe aberrations of the skull greatly degraded imaging resolution and contrast (Fig. 5b). Taking advantage of MeNet-AO, we achieved simultaneous high-resolution *in vivo* imaging of microglial structure and calcium dynamics in CX3CR1-tdTomato-GCaMP7s mice, in which microglia were dually labeled with tdTomato and GCaMP7s.

**Fig. 5.**
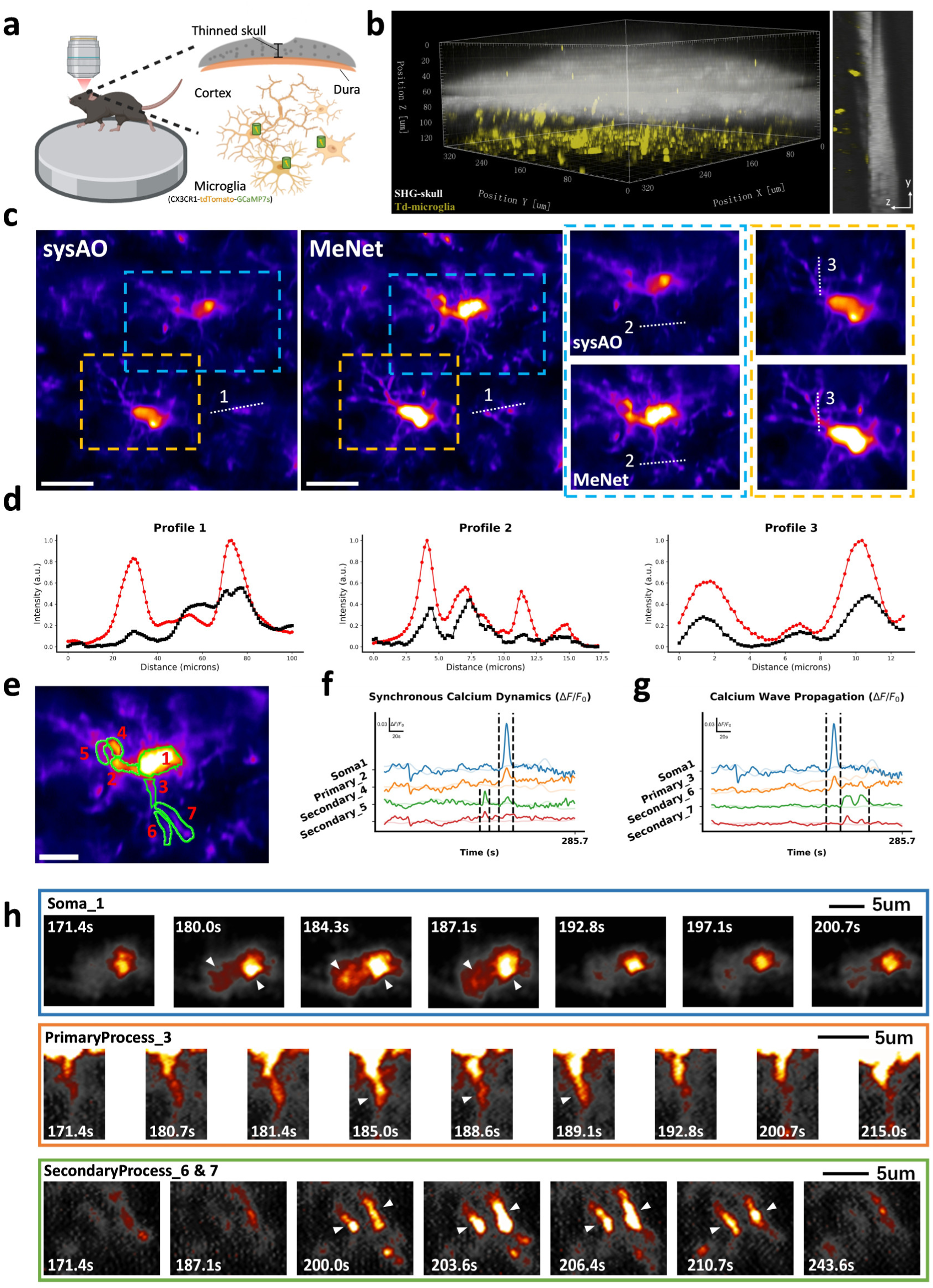
MeNet-AO enables high-resolution structural and functional microglial imaging through thinned-skull windows. **(a)** Schematic of microglial imaging in the cortex of CX3CR1-tdTomato-GCaMP7s mice through a thinned-skull cranial window. **(b)** 3D reconstruction (350×350×130 𝜇𝑚^ଷ^ volume, 5μm z-step) of the thinned-skull cranial window. Gray: second harmonic generation (SHG) signals of skull; Yellow: tdTomato-labeled microglia. **(c)** Representative microglia images acquired at 30 μm below the dura, with system AO (left) and MeNet-AO (right) correction. The images are displayed as average-intensity projections of 400 frames (1.4 Hz, tdTomato channel) for visualization. MeNet-AO correction was performed prior to acquiring the image sequence. Zoomed-in views correspond to the regions marked by the orange and blue dashed boxes. **(d)** Intensity profiles along the dashed lines 1-3 in (c), with system AO (black) and MeNet-AO (red) correction. For direct comparison, the MeNet-AO images were rigidly registered to the system AO reference. **(e)** Structural segmentation of a representative microglia within the region of interest (ROI) outlined in (c). **(f)** The calcium traces (Δ𝐹/𝐹_0_, smoothed) from the soma and processes in ROIs #1/2/4/5 show synchronous calcium dynamics. **(g)** The calcium traces (Δ𝐹/𝐹_0_, smoothed) from the soma and processes in ROIs #1/3/6/7 show a sequential calcium propagation pattern from the soma (#1) and primary process (#3) to secondary processes (#6/7). **(h)** Time-resolved calcium dynamics revealed by MeNet-AO two-photon microscopy, showing delayed activation in secondary processes (ROIs #6/7) following soma (ROI #1) and primary process (ROI #3). Frames marked by white triangular arrowheads indicate calcium activity. Scale bar, 20 μm (c), 10 μm (e).

First, we imaged microglia located 30 μm below the dura. Even at this shallow depth, skull-induced aberrations prevented clear visualization of complete microglial morphology (Fig. 5c). Given the inherently weak GCaMP7s signals in microglia, structural imaging relied on brighter tdTomato fluorescence. However, skull aberrations degraded tdTomato intensity below the threshold required for high-quality spot patterns essential for DWS-AO (Supplementary Fig. 19). In contrast, MeNet-AO, using the same tdTomato signals, enabled robust aberration measurement and correction. As can be seen, with MeNet-AO correction, fine microglial processes can be clearly resolved across the field of view (FOV) (Fig. 5c). Quantitative analysis showed that MeNet-AO correction increased the signal intensity of microglial processes by two- to fivefold (Fig. 5d).

Next, we performed functional calcium imaging of microglia. Unlike neurons, microglial calcium transients typically exhibit lower amplitudes, occur less frequently, and are predominantly confined to processes, making them more challenging to detect^73^. Time-lapse imaging was performed at 1.4 Hz for 400 frames under both AO-corrected and uncorrected conditions. By analyzing the AO-corrected data, we identified two distinct calcium signaling patterns in microglia. After segmenting microglial soma and processes by branch orders based on tdTomato signals, we observed both synchronized and propagating calcium transients. As shown in Fig. 5e&f, somatic calcium transients were accompanied by highly synchronized calcium transients in primary process #2 and secondary processes #4/5, which is consistent with previous studies (Fig. 5f, dark traces: MeNet-AO corrected; light traces: uncorrected). Meanwhile, secondary processes #6/7 displayed delayed calcium transients initiating shortly after the soma and primary process #3, suggesting a propagation pathway (Fig. 5g). The zoomed-in time-lapse image sequences highlight three representative calcium spark events originating from the soma, primary process #3, and secondary processes #6/7, with each exhibiting a distinct sequential activation pattern in their calcium responses (Fig. 5h). Notably, secondary processes #6/7 were barely visible without AO correction (Fig. 5c&d), underscoring the critical role of AO in resolving reliable microglial calcium dynamics (Supplementary Fig. 20a-c). This exhibits that calcium waves propagate selectively along specific processes rather than uniformly, revealing spatiotemporal heterogeneity in calcium dynamics within individual microglia (Supplementary Video). In another microglia (Supplementary Fig. 20d), AO correction enabled detection of a process-specific calcium transient with two consecutive peaks in otherwise undetectable structures (Supplementary Fig. 20d-f, process #11/12/13). These results demonstrate MeNet-AO’s capacity to significantly increase detection likelihood and accuracy of microglial calcium transients, providing a powerful tool for *in vivo* functional study of microglia.

Collectively, MeNet-AO can successfully measure and correct aberrations even under strong scattering and low signal-to-noise conditions, which is challenging for DWS-AO. This minimally invasive microscopy enables subcellular-resolution functional calcium imaging of microglia via thinned-skull windows, opening avenues for high-fidelity analysis of neuron-glia interactions within native microenvironments.

## Discussion

The integration of AO with deep learning has significantly advanced intravital microscopy^8,42–44^, expanding opportunities for diffraction-limited imaging in scattering tissues that pose longstanding challenges. Our MeNet-AO framework addresses two fundamental limitations of conventional AO systems: 1) guide-star dependency in wavefront sensing, and 2) temporal constraints in iterative correction approaches. Compared to conventional sensorless modal AO approaches requiring additional measurements that scale linearly with Zernike mode count^24–29^ (e.g., 2N+1 or 3N acquisitions for N modes), MeNet-AO achieves comprehensive aberration prediction of at least seven Zernike modes (Z_5_-Z_11_) using three modulated image pairs. This is advantageous for correcting high-order aberrations while preserving temporal resolution, which is critical for dynamic intravital imaging. By implementing optimized Zernike mode modulation with a physics-informed multi-encoder architecture, we achieve real-time correction (<5 seconds) with high accuracy. This breakthrough stems from systematic modulation optimization through predictive coupling analysis of Zernike modes, along with noise-robust, structure-independent feature disentanglement via frequency-domain regularization and specialized parallel encoder pathways. The multi-encoder network is specifically crafted to enhance feature extraction for each modulation type, and to prevent the feature mixing and aliasing that occurs when all modulated inputs are stacked and processed by a single encoder. By employing dedicated encoders for each input, we explicitly disentangle the distinct features associated with each modulation (Supplementary Fig. 1c), thereby enhancing cross-modal prediction power. Conceptually, this aligns with the established use of multi-branch architectures in related fields^37,39,74–76^, which are adopted to process modality-specific information before fusion. These improvements in processing speed, generalization capability, and correction accuracy collectively enable real-time aberration correction for intravital imaging, as evidenced by high-resolution functional imaging of neuronal and microglial calcium dynamics in the mouse cortex through open-skull and thinned-skull preparations. To our knowledge, this is the first application of intravital functional imaging to monitor microglial calcium dynamics at subcellular resolution through thinned-skull preparations. This approach offers a powerful platform for high-fidelity investigation of microglial calcium activity while eliminating surgery-induced activation artifacts inherent to open-skull preparations.

While deep learning successfully decouples guide star dependency from correction speed in AO systems, its application to intravital deep tissue imaging presents unique challenges. Unlike single-molecule localization techniques^42,43,77–79^ where PSF-like single-molecule patterns directly encode aberrations or superficial imaging where scattering effects remain manageable, DLAO for intravital deep tissue imaging must confront two compounding challenges: 1) structural complexity that obscures aberration signatures, and 2) signal degradation caused by scattering and large-amplitude tissue aberrations that exacerbates with depth^80^. These challenges render wavefront estimation from single images fundamentally ill-posed. We address this through systematic wavefront modulation that creates additional information channels for aberration isolation. Our optimized modulation strategy synergizes Zernike mode selectivity with noise-robust feature extraction, enabling reliable and high-speed operation in low-signal regimes where traditional guide star-based wavefront sensing fails. Our validation against DWS-AO reveals superior performance of MeNet-AO in detecting weak calcium transients of neurons and microglia through open-skull and thinned-skull cortical imaging. Future efforts could focus on developing a universally effective modulation strategy with improved speed, FOV and imaging depth. In this regard, phase diversity (PD) research offers instructive design principle^61,62,64,81–87^. For example, hybrid modulation schemes that combine different aberration types may shorten the modulation cycle without compromising estimation accuracy. Furthermore, MeNet-AO could be extended by integrating 3D (multi-plane) PD concepts^85^ and more rigorous noise models^87^ derived from PD theory, providing stronger constraints for robust wavefront estimation in complex media. Additionally, spatial encoding architectures combined with pupil-segmentation AO may enable aberration correction across an extended FOV in heterogeneous tissues^79,88^. Furthermore, integrating scattering models into network could enhance performance in optically dense environments, such as intact skull^41^.

While the supervised learning framework of MeNet-AO necessitates balance between physical accuracy and experimental data constraints, employing physical wavefront modulation by deformable mirror rather than simulated aberration models effectively mitigates the mismatch between targeted and actual aberrations. This strategy ensures that the aberration profiles used in training precisely mirror those encountered in actual tissue correction scenarios. Emerging unsupervised methods for jointly estimating aberration-free images and their corresponding aberrations without the need for additional training datasets show great promise for the accessible deployment of AO across various microscopy systems^54,55,64^. However, achieving optimal performance still depends on highly accurate deformable mirror calibration and precise modeling of optical system parameters. Moreover, computational latency remains restrictive for dynamic imaging applications. Future advancements in DLAO may arise from improved feature extraction and aberration estimation models using advanced neural networks^89,90^. Crucially, the development of standardized benchmark datasets for intravital AO imaging would accelerate the evaluation and comparison of emerging methodologies. By addressing the aforementioned challenges, next-generation DLAO systems could achieve real-time, large-FOV correction in complex biological environments characterized by high spatiotemporal heterogeneity, ultimately bridging the resolution gap between intravital and *ex vivo* microscopy.

## Materials and Methods

### Adaptive optics two-photon microscopy

A schematic diagram of the custom AO-TPEFM system is presented in Fig. 2a and Supplementary Fig. 8a. The output beam from a tunable femtosecond laser (Coherent, Chameleon Discovery NX) was expanded by a 1:4 beam expander (Edmund, 45-803 & Thorlabs, AC254-400-B-ML) to fill the aperture of a deformable mirror (DM; Alpao, DM97-15). The reflected beam was de-magnified 1.7 times by a pair of relay lens L3 and L4 (Edmund, 49-368 & 49-363). The beam was further de-magnified 2 times by L5 and L6 (Thorlabs, AC254-150-AB-ML & Edmund, 49-358) to match the entrance aperture of a Resonant-Galvo-Galvo (RGG) scanner system (Vidrio Technologies). Within the RGG scanner system, the resonant scanner and the center of galvo X and galvo Y scanners was conjugated by two lenses of identical focal length (L7 & L8, Thorlabs, LSM04-BB). The beam was then coupled into a scan lens (Thorlabs, SL50-CLS2), tube lens (Thorlabs, TTL200-MP2) and an objective lens (Olympus, 25X, N.A. 1.05, XLPLN25XWMP2). The DM, resonant x scanner, center of galvo X and galvo Y scanners, and the back pupil of the objective lens were all mutually conjugated.

For two-photon fluorescence imaging, the fluorescence signal was reflected by dichroic mirror D2 (Semrock, FF705-Di01-25×36) and was further split by dichroic mirror D4 (Semrock, FF560-Di01-25×36). Each fluorescence channel was detected by a photomultiplier tube (PMT) module (Hamamatsu, H7422PA-40) equipped with proper short-pass and band-pass filters to block the excitation laser and isolate specific emission wavelengths.

For wavefront sensing, the fluorescence signal was transmitted through dichroic mirror D3 (Semrock, FF488-Di01-25×36) and propagated back along the excitation optical path until it was reflected by dichroic mirror D1 (Semrock, FF705-Di01-25×36). The light was then de-magnified by a 2:1 telescope L9 and L10 (L9: f=300 mm & L10: f=150 mm), and directed to the custom-built Shack-Hartman wavefront sensor (SHWS). The SHWS consists of a micro-lens array (SUSS MicroOptics, 18-00197) conjugated with the DM and an electron-multiplying charge-coupled device (EMCCD; Andor iXon Ultra 888). A short-pass (Semrock, FF01-680/SP-25) and a band-pass filter (Semrock, FF01-609/70 or FF01-520/70) were placed before the SHWS to block excitation laser and select the two-photon fluorescence guide star signal. The coordination and control of individual modules were achieved using the ScanImage platform.

### Correction of system aberrations

System-induced aberrations arising from imperfections and misalignments of optical components were measured and corrected prior to experiments using a Zernike-mode-based sensorless AO algorithm^19–21,24^. The fluorescence intensity of rhodamine 6G was used as the optimization metric. The first 21 Zernike modes (excluding tip, tilt, and defocus) were optimized iteratively to maximize the fluorescence signal. For each Zernike mode, seven discrete aberration levels were applied via the DM, and the resulting fluorescence intensities were fitted with a Gaussian curve to identify the optimal correction value. This system aberration correction procedure was routinely performed prior to subsequent imaging experiments.

### Calibration of the SHWS

The SHWS was calibrated with the DM following a previously reported procedure^19–21^. In brief, a fluorescent guide star was generated by two-photon excitation of rhodamine 6G solution. The guide star signal was de-scanned, reflected by the DM, and directed to the SHWS for wavefront sensing. Then, the first 65 Zernike modes were sequentially applied to the DM, and the resulting spot shifts on the SHWS were recorded to form the influence matrix 𝑀_𝑠𝑧_. Each row of the influence matrix represents the corresponding spot shifts of a given standard Zernike mode. The relationship between the wavefront deformation (Δ𝑍_𝐷𝑀_) and the corresponding spot displacements on the SHWS (Δ𝑆_𝑆𝐻_) is given by: Δ𝑆_𝑆𝐻_ = 𝑀_𝑠𝑧_ ∗ Δ𝑍_𝐷𝑀_. After calibration, verification procedures are performed by sequentially applying individual Zernike modes via the DM and comparing them to SHWS measurements (Supplementary Fig. 8b, c)^91^.

### Aberration-related feature extraction with regularization

The aberration-related feature extraction with regularization is a key step for aberration estimation via MeNet-AO method. Given an object 𝑜 and the Point-Spread-Function (PSF) ℎ, we can describe the physical forward imaging model as^92^:

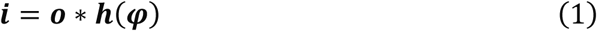

where 𝑖 is the image, and

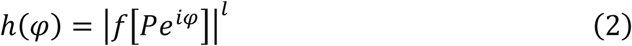

where 𝜑 is the wavefront aberration described as Zernike polynomials

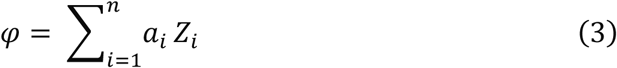

Here, 𝑓[·] is the Fourier transform function, 𝑃 is the Pupil function, and l, which is related to the imaging modality, was set to 4 for two-photon microscopy in our study. 𝑍_𝑖_ is the i-th (Noll index) Zernike mode and 𝑎_𝑖_ represents its corresponding amplitude^93–94^.

For a single image, the presence of two unknown elements, 𝑜 and, makes it an ill-posed problem to estimate the aberrated PSF directly. To address this, we modulated the raw image with pairs of opposite Zernike modes:

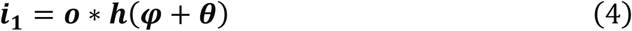

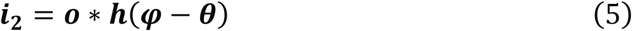

To suppress the influence of sample structural variations, aberration-related features were extracted from each pair of modulated images.

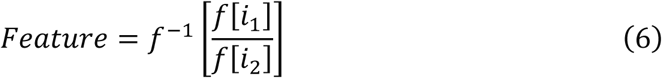

Furthermore, to enhance the robustness of aberration feature extraction against noise, we introduced a regularization strategy (Supplementary Note).

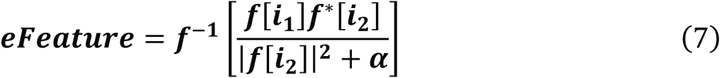

Where **𝑒𝐹𝑒𝑎𝑡𝑢𝑟𝑒** represents enhanced feature, 𝑓^∗^ is the complex conjugate operation, 𝛼 is the regularization value.

During implementation, we computed and generated a pair of features by swapping the order of the image pair (**𝑖_1_**, **𝑖_2_**), and cropped the central 32×32-pixels region as the final input to the network. To address inter–frame motion and ensure robust feature extraction, a shift–correction strategy was applied in which the image region with the highest signal content was first identified, and a fixed-size patch centered on that region was then cropped (Supplementary Note 4). For application of the MeNet-AO model with regularization, the regularization value was set to 1e2 based on optimization results (Supplementary Fig. 9 & 10).

### Simulated aberrated image dataset generation

To determine the optimal combination of modulation modes for MeNet-AO, we created a simulated aberrated image dataset by introducing simulated aberrations into real fluorescence images. Aberration-free slice images containing GCaMP-labeled neurons and DAPI-stained nuclei were acquired using our AO-TPEFM with system aberrations corrected. Based on a physical forward imaging model, simulated aberrations and noise were added to these experimental images to generate aberrated images. In addition, modulated image pairs were generated using predefined modulation modes (Supplementary Fig. 2).

Different strategies were employed to generate random aberrations depending on the specific evaluation objective. To investigate cross-modulation correlations, single-mode aberrations were randomly sampled from the range of -0.5 μm to 0.5 μm, and each was further modulated by a pair of opposite Zernike modes. To generate diverse initial aberration combinations, simulated Zernike coefficients were produced using a semi-random sampling approach. Each mode (Z₅–Z₁₁, Noll index, representing primary aberrations excluding tip, tilt, and defocus) was independently sampled from a normal distribution with a zero mean and a standard deviation of 0.3 μm. For each mode, there was a 30% probability of assigning a non-zero value. Aberration combinations with a total wavefront RMS less than 0.6 μm were retained for further modulation, in which several pairs of Zernike modes were applied sequentially. The resulting modulated aberrated image pairs were fed into the feature extraction pipeline described earlier. The extracted features and their corresponding degraded images served as input-target pairs for supervised training. To train the model, 1,000 samples were generated for single-mode aberrations and 5,000 samples for multi-mode aberrations.

### Experimental aberrated image dataset acquisition

A three-color labeled brain slice with DAPI-labled nuclei, GFP-labeled microglia, and tdTomato-labeled neurons was used to construct the experimental aberrated image dataset. For each region of interest (ROI), the initial aberration modes, generated following the rules described in the section “Simulated Aberrated Image Dataset Generation”, and the corresponding modulation modes were sequentially applied to the image via the DM. Two-photon fluorescence images were acquired at a resolution of 512×512 pixels over a 100 μm×100 μm FOV at 1.22 Hz. The collected image pairs were used to extract aberration-related features, while the initially loaded aberration data served as ground truth for supervised training of MeNet.

To assess the impact of signal-to-noise ratio (SNR) on model performance, 1,000 samples were collected under varying excitation laser powers. In addition, 5,000 samples incorporating all primary aberration modes were acquired from five ROIs to train the model for subsequent *in vivo* imaging applications.

A summary of the key aberration parameters for all datasets presented in the figures is provided in Supplementary Table 1.

### MeNet architecture and network training

MeNet-AO receives aberration-related inputs generated through phase modulation with different Zernike modes and is designed to extract hierarchical feature representations, which are subsequently integrated to reconstruct the final wavefront in terms of Zernike coefficients. The network architecture consists of three parallel feature encoders, followed by a feature integration module.

Specifically, each encoder begins with a convolutional layer, followed by batch normalization and activation via a parametric rectified linear unit (PReLU). To capture spatial information within the small cropped regions, we used a relatively large kernel size – 7×7 pixels in the first layer and 5×5 pixels in the second. This is followed by a residual group composed of four stacked residual blocks, with the number of channels increasing from 16 to 256, designed to extract rich and diverse features^95^. A final convolutional layer follows the residual group. Since aberration features modulated by different Zernike modes exhibit distinct characteristics, we constructed a multi-encoder structure, in which each encoder (with the same architecture) processes input corresponding to a specific modulation. The extracted features are then adaptively integrated by a feature integration module, which also includes a stacked residual group used to predict the final wavefront. Finally, two fully connected layers are used to regress the amplitudes of the Zernike modes. For the single-encoder setting, to match the parameter count of MeNet, the residual group is expanded to five stacked residual blocks, with the number of channels increasing from 16 to 256, directly followed by one additional residual block with 256 channels. This adjustment resulted in comparable parameter counts (single encoder 4.62M vs. MeNet-AO 4.50M), demonstrating that the superior performance of MeNet-AO is attributable to its multi-encoder architecture, not simply to model size.

For a direct benchmark, we reimplemented the MLAO framework by strictly following its original CNN architecture^63^. Our implementation achieved comparable performance (Supplementary Fig. 6b), thus ensuring a valid comparison.

In this work, the MeNet networks were trained on workstations equipped with NVIDIA RTX 3090Ti or RTX 4090 GPUs, using Python 3.9 and TensorFlow 2.6.0. During training, the dataset was split into a 9:1 ratio for training and validation. The model was optimized using the Adam optimizer with an initial learning rate of 6×10^-4^, which decayed by a factor of 0.5 every 10,000 iterations. Training was performed for 1,000 steps per epoch with a batch size of 64. Mean squared error (MSE) was adopted as the loss function. The number of epochs was adjusted according to the complexity of the testing task, typically set to 100 or 200. Default architectures and hyperparameters were maintained across methods (SingleEncoder, MeNet-AO, MLAO) and all models were trained/tested on identical datasets.

### MeNet-AO and DWS-AO correction

#### MeNet-AO

In MeNet-AO correction, three predefined Zernike modes—oblique astigmatism, vertical astigmatism, and primary spherical aberration—were applied to the initial image via the DM. Images were acquired at a resolution of 512×512 pixels over a 100 μm×100 μm FOV at 1.22 Hz. The resulting image pairs were saved and subsequently loaded into a Python-based pipeline. Aberration-related features were then extracted as described above and input into the trained MeNet model. The correction modes predicted by MeNet-AO were applied to the DM for wavefront compensation.

#### DWS-AO

Direct wavefront sensing (DWS) AO was performed according to a previously established protocol^19–21^. After SHWS calibration, the spot pattern 𝑆_𝑟𝑒𝑓_ obtained with system correction was saved as a reference for subsequent sample aberration measurements. In this study, neuronal or microglial fluorescence signals were used as guide stars to capture the spot pattern 𝑆 in zebrafish and mouse brains.

Then the corrected modes could be computed by: 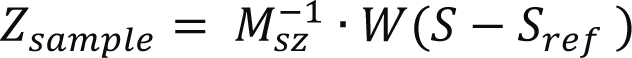, where 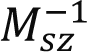 denotes the generalized inverse matrix and ܹ is a weighted matrix determined by the signal-to-background ratio of each spot on the SHWS. Finally, the full AO correction is applied as the combination of system and tissue aberrations.

### Animal preparation

Zebrafish were maintained according to standard protocol^96^. *Tg(Neurod:GFP)* and *Tg*(*Neurod:GFP; flk1:DsRedx*) lines were used in this study. Multiple transgenic mouse lines were utilized in this study. Neuronal imaging was performed using Thy1-GCaMP6f mice. For direct wavefront sensing experiments, stable guide stars were generated by injecting AAV2/9-hSyn-jGCaMP8s-P2A-NLS-Dendra2 into targeted brain regions of C57BL/6J mice, resulting in bright nuclear-localized neuronal fluorescence. Intravital imaging was conducted three weeks after viral injection. For microglial studies, CX3CR1-tdTomato-GCaMP7s mice were employed to simultaneously monitor morphology and calcium dynamics in microglia. All animal experimental procedures were conducted in accordance with animal care guidelines approved by the Southern University of Science and Technology Animal Care and Use Committee. Before imaging experiments, animals were housed on a 12-hour light-dark cycle with food and water ad libitum.

### Fixed mouse brain slices preparation

Brain slices were prepared from Thy1-GCaMP6f mice expressing GCaMP6f in neurons, and CX3CR1^GFP/+^ mice expressing GFP in microglia. To test model generalization (Fig. 2e-h, Supplementary Fig. 12), brain slices with triple-color labeling were prepared. Specifically, AAV-hSyn-tdTomato was injected into the primary visual cortex (V1) of CX3CR1^GFP/+^ mice to label neurons (tdTomato) and microglia (GFP) simultaneously. After 3-4 weeks of viral expression, mice were transcardially perfused with phosphate-buffered saline (PBS), followed by 4% paraformaldehyde (PFA). Brains were post-fixed, cryoprotected in 30% sucrose, and coronally sectioned at 20 μm thickness. Tissue sections were stained with DAPI for nuclei visualization and mounted for imaging. Experimental image datasets were collected from sections in which microglia (GFP), neurons (tdTomato), and nuclei (DAPI) were simultaneously visible.

### *In vivo* zebrafish imaging

The 4-dpf larvae were anesthetized in 0.02% Tricaine prepared in E2 egg water, then mounted in 1% low-melting-point agarose within 35 mm glass-bottom confocal dishes (Biosharp, BS-15-GJM). For imaging, larvae were oriented either laterally or dorsally and placed in direct contact with the glass to ensure stable positioning on the dish.

### Open-skull cranial window surgery for mouse cortical imaging

For neuronal imaging in mouse cortex, open-skull cranial window surgery was performed following established protocols^97^. Mice (8–12 weeks old) were anesthetized with 4% isoflurane for induction and maintained at 1–2% isoflurane throughout the procedure. The scalp was retracted, and the periosteum was carefully removed from the exposed skull. A circular craniotomy (3–4 mm in diameter) was performed using a high-speed dental drill at a site located 3 mm lateral to the midline and 4 mm posterior to bregma. The bone flap was gently lifted, ensuring the dura mater remained intact. A 4 mm circular cover glass was affixed above the dura using cyanoacrylate adhesive. The surrounding skull was coated with a thin layer of dental cement, and a custom-designed head plate with a 9 mm central aperture was aligned concentrically with the cover glass and fixed to the skull. Additional dental cement was applied around the head plate to seal the surgical site and ensure mechanical stability.

### Optical clearing and thinned-skull window surgery

For microglial imaging, we implemented an optically cleared thinned-skull cranial window to minimize surgery-induced activation. The procedure for optical clearing of the skull window followed a previously published protocol with minor modifications^19,72^. Two reagents were used: S1, a saturated supernatant solution consisting of 75% (v/v) ethanol and urea at room temperature; and S2, a sodium-based solution prepared by mixing 0.7 M NaOH with an acid at a volume-to-mass ratio of 24:5.

During surgery, mice were anesthetized with isoflurane as described above. Before applying the clearing reagents, the compact outer layer of the skull was carefully thinned under a surgical microscope using a high-speed micro-drill. This polishing step reduced the skull thickness to approximately 40–60 μm while preserving the inner trabecular layer. Care was taken to avoid damaging the underlying dura mater or causing bleeding. This procedure improved reagent penetration and enhanced optical clarity of the skull for subsequent imaging. A custom-designed head plate was affixed to the skull and secured with dental cement. Reagent S1 was applied to the exposed skull surface for approximately 20 minutes, during which a clean cotton swab was used to gently rub the surface and accelerate the clearing process. After removing S1, reagent S2 was applied to the skull surface for 5 minutes. Following this, a fresh drop of S2 was reapplied to the skull for refractive index matching. Finally, a 5-mm-diameter cover glass was placed over the treated area and sealed along the edges with cyanoacrylate adhesive.

### *In vivo* two-photon imaging of mouse brain

During intravital cortical imaging, mice were anesthetized with isoflurane and head-fixed using a custom-designed mounting stage. The cranial window was aligned perpendicular to the optical axis of the objective. The femtosecond laser was tuned to 920 nm, and post-objective excitation power ranged from 20 to 50 mW depending on imaging depth.

For two-photon calcium imaging of neurons, mice were maintained under light anesthesia (0.5% isoflurane). Imaging was performed in resonant–galvo scanning mode at a frame rate of 30 or 15 Hz, with a resolution of either 512 × 512 or 1024 × 1024 pixels. Visual stimuli were presented to the eye contralateral to the imaged hemisphere using a liquid crystal display (LCD). The stimuli consisted of full-field, square-wave drifting gratings in eight directions (100% contrast, 0.035 cycles/degree, 2 Hz), with each direction presented for 2 seconds and interleaved by a 1-second gray blank screen.

For two-photon imaging of microglia, isoflurane was administered at an increased flow rate to maintain a concentration between 3% and 4%^73^. Imaging was performed in galvo-galvo scanning mode at a frame rate of 1.4 Hz with a resolution of 512 × 512 pixels.

### Image processing and analysis

#### Structural imaging analysis

All images were processed with ImageJ and Python. The raw images were denoised using the Smooth function in ImageJ. The image pairs with system AO and full AO correction were acquired and processed using the same parameters and displayed with consistent contrast. For comparison, the image pairs were registered using the *Linear Stack Alignment with SIFT* plugin in Fiji software^98^. The frequency spectrum analysis of the images was performed using a custom Python script. For volumetric imaging of zebrafish neurons and the mouse skull, 3D reconstruction and visualization were performed using Imaris 9.0 (Oxford Instruments).

#### Neuronal functional imaging analysis

Time-lapse recordings of neuronal calcium activity during visual stimulation were motion-corrected, background-subtracted, and segmented using the Python implementation of Suite2p^99^. Calcium traces extracted by Suite2p were neuropil-corrected and subsequently imported into a custom Python analysis program. The precise timing of stimulus onset and interstimulus intervals was determined based on background light leakage signals at signal-free regions. Calcium responses were quantified as Δ𝐹/𝐹_0_, where 𝐹_0_ represents the baseline fluorescence signal during the non-stimulation period. ROIs corresponding to direction/orientation-selective responses were manually selected from the automatically generated ROI set. To quantify the AO-induced enhancement in orientation selectivity^100,101^, multiple ROIs were manually defined and the motion-corrected calcium traces were extracted using the Multi-Measure tool in Fiji.

#### Microglial functional imaging analysis

Microglial calcium activity was analyzed in Fiji and Python, with reference to prior studies^73^. An average intensity image of the tdTomato channel, which reflects cell morphology, was generated for manual ROI selection. Microglial somata, primary processes, and secondary processes were identified based on their size and morphology. Using the manually selected ROIs, the Multi-Measure tool in ImageJ was used to extract mean fluorescence intensity values, which were exported and loaded into a custom-built Python program for conversion into Δ𝐹/𝐹_0_. The baseline fluorescence (𝐹_0_) was calculated by smoothing the signal over non-responsive time windows across all frames. To improve the visualization of weak GCaMP signals in microglia, the raw data were enhanced using SRDTrans^89^ for display purposes only.

#### SNR analysis

To quantitatively evaluate the model’s performance on data with varying noise levels, peak signal-to-noise ratio (PSNR) was used as the evaluation metric.

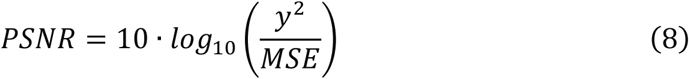

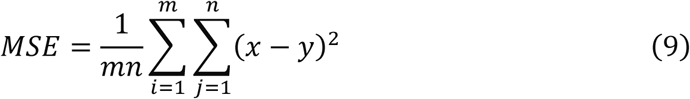

Here, y represents the reference image acquired under the highest excitation laser power, and x denotes the noisy image obtained with lower excitation laser power.

## Data and Code availability

The data and source code for MeNet-AO will be released after review.

## Acknowledgements

We thank Prof. Yong U. Liu at South China University of Technology for providing the CX3CR1-tdTomato-GCaMP7s mice, and thank Prof. Zilong Wen at Southern University of Science and Technology for sharing the *Tg(Neurod:GFP; flk1:DsRedx)* zebrafish. This work was supported by the National Natural Science Foundation of China (32192400), Shenzhen Medical Research Fund (B2301004), the Guangdong Basic and Applied Basic Research Foundation (2024A1515012414), the National Key R&D Program of China (2024YFA1804000, 2023YFA1800100), the National Natural Science Foundation of China (32101211), and the Guangdong Innovative and Entrepreneurial Research Team Program 2021ZT09Y104.

## Author contributions

X.C. and S.H. conceived the project. S.H. supervised the work. X.C. and S.H. developed the MeNet-AO algorithm, designed the experimental schemes, analyzed the data, and prepared the figures. L.L., X.C., and S.H. built the AO two-photon microscope system. B.W. and L.L. performed the mouse brain surgeries. X.C., B.W., and L.L. conducted the mouse brain imaging experiments and acquired the corresponding data. X.C. and Z.S. performed zebrafish imaging and data acquisition. X.C. and S.H. wrote the manuscript with input from all other authors.

## Competing interests

All authors declare no competing interests.

## Notes

### Competing Interest Statement

The authors have declared no competing interest.

